# MegaMASLD: An interactive platform for exploring stratified transcriptomic signatures in MASLD progression

**DOI:** 10.1101/2024.07.21.603199

**Authors:** Hong Sheng Cheng, Damien Chua, Sook Teng Chan, Kuo Chao Yew, Sunny Hei Wong, Nguan Soon Tan

## Abstract

Liver transcriptomic data from patients with metabolic dysfunction-associated steatotic liver disease (MASLD) offers valuable resource for deciphering pathogenic molecular drivers. Here, we performed a Mega-analysis of MASLD Liver Transcriptomes (MegaMASLD) which reanalysed raw RNAseq data of over 800 livers in a standardized and integrative manner, aiming to unravel druggable molecular events in MASLD. Our analysis revealed a progressive transcriptomic shift predominantly associated with immunopathologies during MASLD progression. The differential transcriptomes produced a MASLD gene signature useful for quantitative assessment of MASLD severity but failed to faithfully recapitulate the exact histological staging. Instead, a histologic-independent unsupervised clustering analysis predicted a high-risk group prone to develop metabolic dysfunction-associated steatohepatitis (MASH), characterized by aberrant changes in humoral immune response and antibody repertoires. These findings were supported by another histologic-independent pseudotime analysis, which also identified several potentially targetable molecular switches, including FGFR, PDGFR, PAK, PRKG1 and CAMK kinase families, activated at various transitory phases of MASLD. The robust analysis has enabled risk stratification and deepened our understanding of the dynamic molecular events driving MASLD, thereby offering new options to enhance precision medicine of MASLD. An online web tool featuring MegaMASLD is available at https://bioanalytics-hs.shinyapps.io/MegaMASLD/.

## Introduction

Metabolic dysfunction-associated steatotic liver disease (MASLD), formerly known as non-alcoholic fatty liver disease (NAFLD), is rapidly emerging as a leading cause of liver-related morbidity and mortality worldwide. The global prevalence of MASLD is estimated to have exceeded 30% in adults, and with highest rates in Latin America (44%), followed by Middle East and North Africa (37%), and South Asia and South-East Asia (33%)^1,2^. The ever-growing prevalence warrants an immediate attention to alleviate the healthcare burden caused by the chronic liver disease. The recent approval of resmetirom, a thyroid hormone receptor β agonist and the first drug that resolves liver fibrosis^3^, by the United States Food and Drug Administration (FDA) for the treatment of metabolic dysfunction-associated steatohepatitis (MASH) highlights a remarkable milestone in curbing the chronic liver disease. In the phase III trial, approximately 30% of the MASH patients (9.7% in the placebo group) had no worsening or improved fibrosis after a one-year course of daily resmetirom^3^. It is, however, worth noting that majority of the patients continued to have progressive MASH and the response rate after placebo adjustment was a 16.4-20.7% improvement in MASH resolution and a 10.2-11.8% improvement in fibrosis^4^. These findings underscore the need for longer treatment duration to observe greater clinical benefits, as well as the development of patient stratification strategy or adjunct therapeutic due to the complex pathogenesis of MASLD.

The analysis of the multiomic landscape forms the foundation of modern precision medicine, driving continuous efforts to identify better molecular markers and novel drug targets to improve MASLD patient outcomes^5,6^. A recent genome-(GWAS) and phenome-wide association study (PheWAS) profiled the genetic architecture and phenotypic data, revealing 17 gene variants that increase the predisposition to MASLD through seven metabolic-related mechanisms^7^. Notably, some of these loci are in well-known susceptibility genes such as *PNPLA3*, *TM6SF2*, *GCKR* and *MBOAT7*^8–10^. However, despite these discoveries, genetic risk factors have not been integrated into MASLD screening and patient stratification likely due to variable genetic contribution and multifactorial nature of the disease. Conversely, transcriptomic profiling has gained traction in MASLD research, providing insights into key molecular events that drive the transition from healthy livers to various disease stages. Differential transcriptomic analyses enable researchers to identify molecular signatures indicative of MASLD exacerbation and resolution^11–14^. Integrating biopsy-derived liver transcriptomes from different MASLD stages, patient demographics and independent research groups can generate robust findings with broad implications for precision treatment of MASLD. However, existing integrative analysis often use a meta-analysis approach that relies on aggregated data, *e.g.* the differentially expressed genes (DEGs) between different stages^11,15^. While convenient, the use of aggregated data in these meta-analyses are potentially interfered by variations in experimental designs, patient populations, and data processing methods across studies, thus obscuring subtle differences in gene expression patterns that are critical for understanding the pathogenesis of MASLD.

To overcome these limitations, a mega-analysis approach which reprocesses and harmonizes the individual-level raw sequencing data from heterogenous sources, not only provide a stronger statistical power, but also enable a more accurate understanding of the molecular events underlying MASLD^16^. When properly executed, a mega-analysis of large transcriptomic data can even reveal the temporal dynamics of MASLD development, paving the groundwork for targeted interventions to intercept and potentially reverse disease progression at its earliest stages. Clearly, such systematically integrated MASLD liver transcriptomic dataset is undoubtedly an invaluable resource for data exploration. By developing the integrative analysis into a user-friendly tool, it is possible to dramatically amplify its outreach and clinical impact. For instance, several interactive tools such as cBioPortal^17^ and Gene Expression Profiling Interactive Analysis 2 (GEPIA2)^18^, are becoming the workhorses driving data exploration in precision oncology. In contrast, similar tools for MASLD are conspicuously limited^19,20^. Despite growing amount of transcriptomic data in MASLD, there is still a lack of user-friendly web-based tools that enable molecular biologists and clinicians, especially those with little or no computational background, to tap into these rich resources for diagnostic and drug development.

This study aims to tackle two important hurdles in MASLD research, the integration of diverse transcriptomic datasets and the accessibility of advanced user-friendly, MASLD-centric web-based analytical tools. Firstly, using an integrative approach, we performed a mega-analysis using publicly accessible raw liver RNAseq data from healthy individuals and those with MASLD. This analysis aims to unravel key molecular events that occur during the temporal dynamics of MASLD pathogenesis. Secondly, we developed Mega-analysis of MASLD Liver Transcriptomes (MegaMASLD), an interactive R Shiny tool offering user-friendly analytical and visualization utilities, including differential expression analysis, gender analysis, correlation studies, transcriptomic-guided staging and pseudotime analysis (**Figure 1A**). By providing these applications, MegaMASLD aims to enhance the accessibility of complex transcriptomic data, thus delivering broad implications in the precision medicine of MASLD.

**Figure 1.**
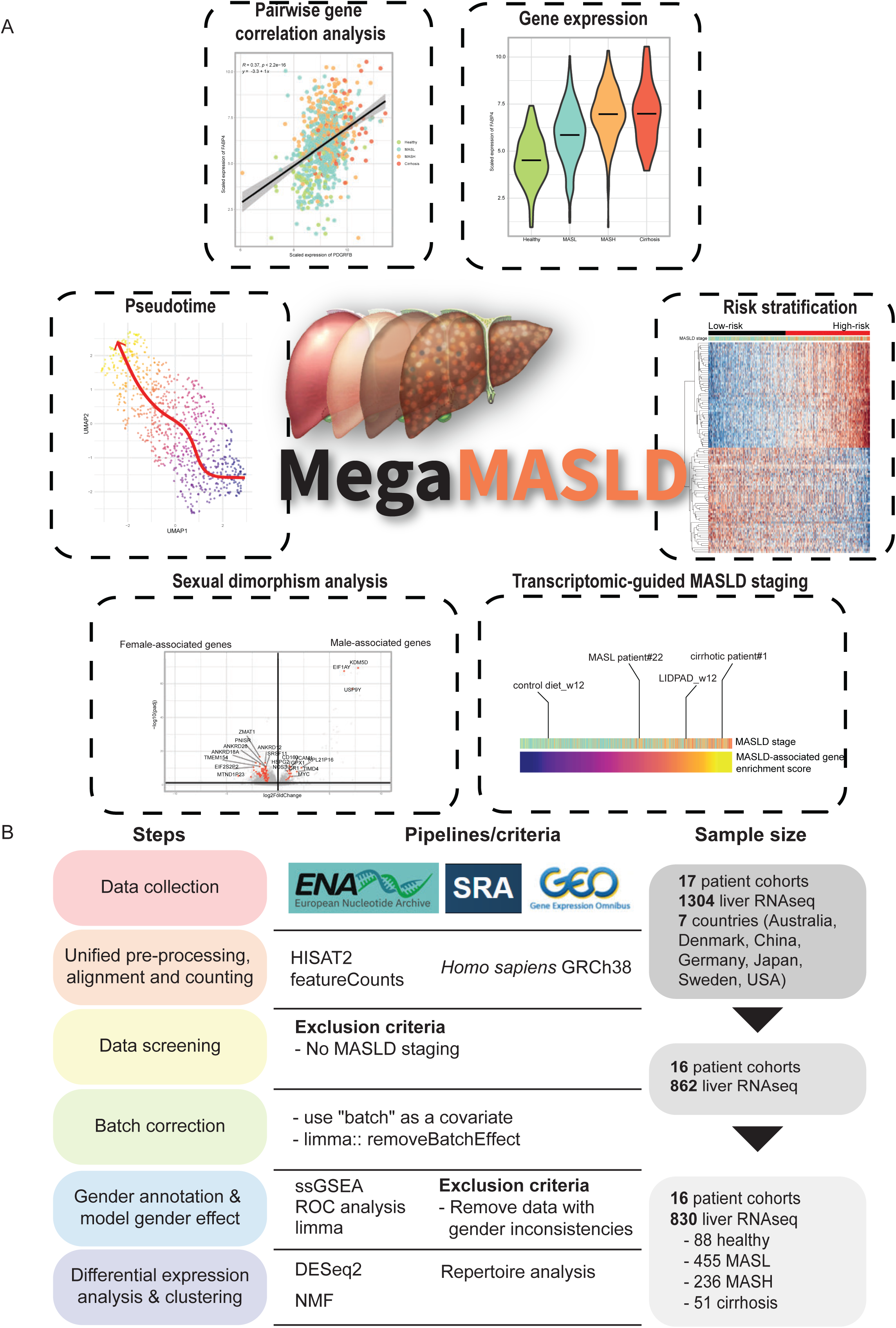
MegaMASLD: An interactive webtool for exploratory analysis of MASLD liver transcriptomes. **A.** The analytical utilities available in MegaMASLD. **B.** A schematic diagram illustrating the collection, processing and data integration of liver transcriptomes from MASLD patients.

## Materials and Methods

### MASLD liver transcriptomes and data pre-processing

Publicly accessible datasets containing human liver transcriptomes of healthy and MASLD individuals were obtained from the Gene Expression Omnibus (GEO) and their raw sequencing data were downloaded from Sequence Read Archive (SRA) and European Nucleotide Archive (ENA) in July 2023. A total of 1304 liver transcriptomes from 17 independent datasets were identified (**Supplementary Data 1**). We employed a unified pre-processing pipeline for genome alignment and gene counting to minimize variability due to methodological inconsistencies (**Figure 1B**). All raw RNAseq reads were mapped to the human genome (GRCh38.p14) using Hisat2^21^, followed by gene counting using featureCount^22^ and human GTF files from Ensembl to obtain the gene count matrices.

### Clinical staging based on liver histology scores

Some liver transcriptomes have liver histology scores, namely NAFLD Activity Score (NAS) and fibrosis staging, but not the clinical staging. To classify the transcriptomes into either healthy, MASL, MASH or cirrhosis, we used the criteria outlined in Table S1^23^.

### Batch correction and data integration

To minimize batch effect due to sequencing runs, service providers and sequencers, different GEO datasets were considered as separate batches and included as a covariate in the statistical design during differential expression analysis to adjust for the batch effect. For visualization, the modelled batch effect was eliminated from normalized and variance stabilizing transformed (vst) gene counts using the *removebatcheffect* function in LIMMA^24^. Principal component analysis was used to inspect the effectiveness of batch correction and presence of other covariance in the integrated datasets.

### Differential expression analysis, functional enrichment analysis and gene-gene interactions

Differential expression analysis between MASLD stages, genders and clusters was performed using DESeq2^25^. All differential expression analyses were done with batch and gender correction, except for gender analysis in which only batch correction was performed to unravel disparities in the liver transcriptomes between males and females. Unless mentioned otherwise, genes with FDR < 0.05, log2 fold change > ±1 and baseMean>100 were considered as DEGs. Functional enrichment analysis was performed using ViSEAGO^26^ or Gene Set Enrichment Analysis (GSEA)^27^. Enrichment analyses of KEGG, protein kinases and transcription factors were carried out using Enrichr^28^. Gene-gene interaction was analysed using Cytoscape^29^.

### Bulk RNAseq deconvolution

Deconvolution of the bulk liver RNAseq into cellular subpopulations was performed using Cibersortx^30^. Gene signatures of 16 intrahepatic cellular subpopulations, including immune cells, hepatocytes, and stromal cells, were derived from single-cell RNAseq data annotated by Li et al. (2023)^19^ using “Create Signature Matrix” function in Cibersortx. The gene signatures and batch-corrected normalized gene counts of the MASLD liver transcriptomes were loaded to impute cell fractions (permutations=1000).

### Transcriptomic-guided gender annotation

Gender was found to be an important covariate in the integrated liver transcriptomes, but a subset of the data lacks the gender information. To predict the gender of these samples, we curated 39 gender-specific markers (28 male-specific and 11 female-specific; **Supplementary Data 2**) based on the DEGs (FDR < 0.05 and log2 fold change > ±1) of liver transcriptomes with known genders followed by a feature selection algorithm, Boruta, to rank the importance of each DEGs and shortlist the important ones for gender classification^31^. Next, the enrichment scores of these male- or female-specific markers in each liver transcriptome were computed using ssGSEA2^32^. Using transcriptomes with known genders, three binomial logistic regression models was created using enrichment scores of male-, female- or male + female-specific markers as the predictors and the genders as the outcomes. Receiver operating characteristic (ROC) curves constructed to evaluate the most predictive model for gender annotation.

### Human-mouse harmonized MASLD signature and transcriptomic-guided staging

The procedure to establish human-mouse harmonized MASLD signature has been described previously^33^ Briefly, human DEGs which were progressively upregulated with increased diseased severity were subject to Boruta feature selection, resulting in 56 signature genes^31^. Addition manual curation was performed to select genes with a mouse ortholog and showing a consistent expression trend with human hepatic expression profile. This resulted in 39 signature genes, also known as human-mouse harmonized MASLD signature (**Supplementary Data 3**). To demonstrate the utility of the curated signature in transcriptomic guided staging, vst gene counts of human and mouse liver transcriptomes with varying disease severity were merged based on shared genes, followed by ssGSEA to compute an enrichment score for each sample^32^. The enrichment scores were used to rank and map the murine MASLD disease severity relative to the human counterparts.

### Risk stratification using consensus non-negative matrix factorization (cNMF)

To uncover potential hidden clusters or subgroups, we performed a cNMF-based unsupervised clustering, a widely adapted method for stable molecular subtyping of malignancies. Using *NMF* package^34^, the top 1000 most variable genes from the integrated dataset was used for 10 iterations of NMF analysis with varying ranks based on Brunet algorithm. Consensus maps and other metrics like silhouette score, cophenetic correlation coefficient were used to evaluate the robustness of the clusters from cNMF.

### B-cell receptor (BCR) repertoire analysis

The third complementarity determining region (CDR3) regions of BCRs were extracted from the raw sequencing data for the profiling of immunoglobulin heavy and light chain repertoires using MiXCR^35^. BCR repertoire diversity analysis and V(D)J gene usage were carried out using *LymphoSeq*^36^. As the RNAseq of the liver biopsies were not specifically designed for BCR repertoire analysis, the sequencing depth might be underpowered to reliably capture the BCR transcripts. Hence, we included only the samples with more than 100 gene counts and 20 different BCR clonotypes in the analysis. Physicochemical properties of the CDR3 region of the light and heavy chain immunoglobulins were evaluated using *Peptides*^37^.

### Bulk RNAseq pseudotime analysis

To ensure robustness of the pseudotime trajectory inference, potential outliers were identified based on interquartile rule of the principal component (PC) 1 and 2. Liver transcriptomes with PC1 or PC2 less than first quartile - 1.5* interquartile range (IQR) or third quartile + 1.5*IQR, were omitted from the analysis. The remaining dataset was comprised of 783 samples (79 healthy, 441 MASL, 225 MASH and 38 cirrhosis). Assuming that the Euclidean distance between liver transcriptomes in a reduced dimensional space reflects meaningful biological relationship, we applied a Gaussian mixture modelling (GMM) onto the PC1 and PC2 using *mclust* package^38^ which automatically identified two clusters based on the Bayesian information criterion. Based on the GMM clustering information and dimensionality reduction from PCA, we performed a pseudotime trajectory simulation using *slingshot* package^39^ which revealed single trajectory that largely aligns with sequential MASLD progression from healthy-MASL-MASH-cirrhosis. A pseudotime value was assigned to each transcriptome to represent its temporal position along the pseudotime trajectory.

To study the (pseudo)temporal gene expression and identify when the expression trend of a gene surges drastically as the disease progresses, we developed a mathematical algorithm to compute a take-off value of all detectable genes. A take-off value is defined as a pseudotime point when the expression of a gene begins to exhibit a significant continuous upward trend. To elaborate, for each gene, we fitted a locally weighted scatterplot smoothing (loess) model to predict the normalized and transformed expression levels (response variable) based on the pseudotime values (predictor variable) (*vst* ∼ *pseudotime*). We excluded all genes showing reduced expression over pseudotime. Next, we computed the first derivative of the loess model, reflecting the rate of change of expression level over pseudotime. A baseline was determined based on the highest first derivative within the earlier pseudotime values (<1.5) whereas an upward trend was defined as 10 consecutive pseudotime points that show at least 0.1% increment in the first derivatives, respectively. A take-off value is assigned to a gene when the first pseudotime point with a first derivative that is at least 10% higher than the baseline and demonstrates an upward trend was detected.

### Statistical analysis

Unless mentioned otherwise, statistical analysis was performed with two-tailed unpaired T test or one-way ANOVA followed by pairwise comparison using Benjamini-Hochberg adjustment. A p-value of less than 0.05 was considered statistically significant.

### Data and code availability

The GEO accession numbers of the hepatic RNAseq datasets used in this study are GSE105127, GSE107650, GSE115193, GSE126848, GSE130970, GSE147304, GSE160016, GSE162694, GSE167523, GSE173735, GSE174478, GSE185051, GSE192959, GSE193066, GSE193080, GSE207310, and GSE213621. Raw sequencing data were downloaded from SRA and ENA. Liver transcriptomes from LIDPAD mouse models were obtained from GSE159911.

Metadata of the liver transcriptomes and data supporting the key findings are available within the article and its supplementary data files. R scripts of various analyses performed in the study is available at https://github.com/chenghongsheng/MegaMASLD. An interactive web server based on the integrated dataset has also been created and publicly accessible at https://bioanalytics-hs.shinyapps.io/MegaMASLD/.

## Results

### Integrative liver transcriptomes revealed metabolic dysfunction, immunopathologies and gender disparity in MASLD

We set out to elucidate the crucial molecular events driving the progressive MASLD from healthy livers to MASL, MASH and cirrhosis using an integrative transcriptomic approach. A thorough data mining in GEO database identified 20 independent RNAseq datasets derived from human liver biopsies of healthy and MASLD individuals. Of these, 17 datasets, comprising 1304 samples from 7 countries from North America, Europe, Asia, and Australia, had publicly accessible raw sequencing data deposited in either NCBI SRA or ENA. A unified and standardized preprocessing pipeline was applied on all the raw sequencing data to minimize inconsistencies arising from bioinformatic pipeline, such as differences in versions and choices of human reference genomes and gene structure files (*e.g.* GTF and GFF files), as well as software used for trimming, genome mapping, and gene counting (**Figure 1B**). After excluding samples that lacked MASLD staging or had received non-standardized treatment potentially affecting the liver transcriptomes, we integrated and analyzed 862 hepatic RNAseq samples from 16 patient cohorts.

Despite the unified preprocessing method, a noticeable batch effect from different studies and sequencing runs undermined the biological differences due to MASLD (**Figure 2A**). To account for this batch effect, we included “batch” representing different GEO datasets in the statistical design. Intriguingly, PCA after batch correction revealed two clusters accounting for about 17% of the variance, which were found to due to gender differences. This finding suggested that male and female livers exhibited intrinsic transcriptomic variation that could potentially interfere downstream data analysis. Furthermore, some transcriptomes lacked the gender information. To annotate the gender of these samples, we developed a transcriptomic-guided gender prediction tool based on selected gender-specific marker genes from 118 DEGs between male and female livers (**Figure S1A, Supplementary Data 2**). Using the enrichment scores from male-specific genes, we achieved high sensitivity and specificity gender prediction (0.96 and 0.95, respectively) at 95.5% accuracy and 0.961 (0.943-0.977; 95% confidence interval) area under curve (AUC) in a ROC analysis (**Figure S1B**). The gender prediction tool was used to annotate transcriptomes with unknown gender. After omitting samples with inconsistent predicted and reported gender, 830 transcriptomes remained, consisting of 88 health, 455 MASL, 236 MASH and 51 cirrhotic livers (**Figure 1B, Supplementary Data 1**). Adjusting for batch and gender successfully reconciled the liver transcriptomes from diverse sources, showing a 12% variance attributable to MASLD stages (**Figure 2A**).

**Figure 2.**
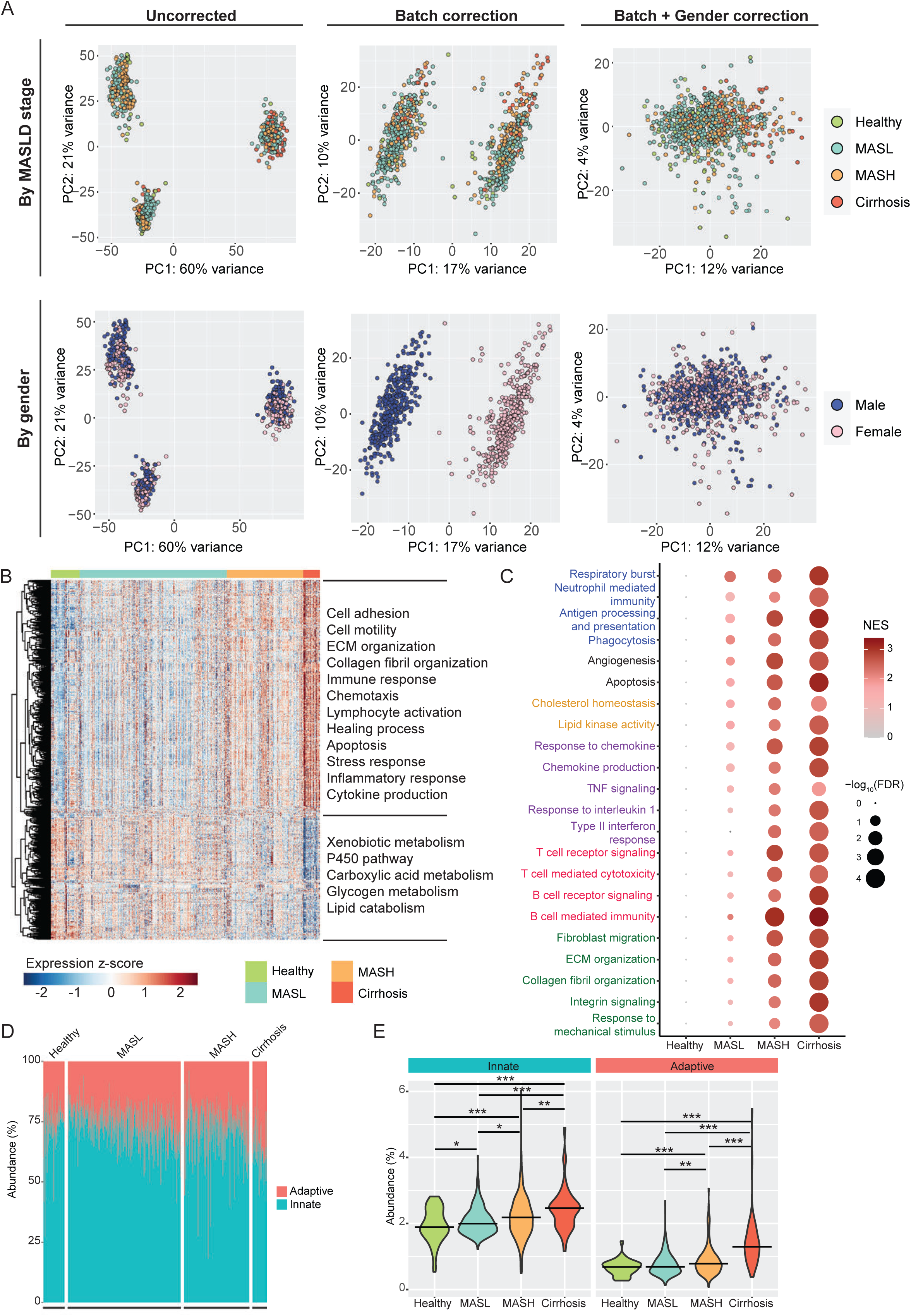
Integrative transcriptomic analysis reveals cumulative immunopathologies in MASLD progression. **A.** Principal component analysis (PCA) highlights batch and gender as the leading covariates explaining substantial variance in the liver transcriptomes from independent datasets. Batch and gender correction enables integration and assessment of transcriptomic shifts due to MASLD progression. **B.** Heatmap of 820 DEGs between MASLD stages and their associated biological functions. **C.** Gene set enrichment analysis (GSEA) of differential transcriptomes between MASL, MASH or Cirrhosis with healthy livers **D-E.** Predicted relative (**D**) and absolute (**E**) abundance of innate and adaptive immune cells at different MASLD stages based on bulk RNAseq deconvolution using Cibersortx. Crossbars indicate median abundances. *p< 0.05, **p< 0.01, ***p < 0.001 (pairwise t test with Benjamini-Hochberg adjustment).

Pairwise differential expression analysis between MASLD stages revealed 820 DEGs, of which 543 were upregulated and 277 were downregulated as the liver disease advanced (**Figure 2B, Supplementary Data 4**). Functionally, the downregulated DEGs were linked to the detoxification activity, carboxylic acid metabolism, glycogen metabolism and lipid catabolism, suggesting a loss of physiological hepatic function and suppressed lipid breakdown which were linked to loss of liver zonation during early steatosis. The upregulated DEGs were over-represented by immune-related genes, demonstrating classical immune pathways like cytokine production, inflammatory response, and chemotaxis. The activation of immune response was accompanied by heightened stress response, apoptosis and healing processes which indicate increased hepatic injury and regeneration in response to inflammatory, oxidative, and metabolic assaults. Some upregulated DEGs were also involved in extracellular matrix (ECM) and collagen organization, highlighting an ongoing tissue remodeling and scar formation within the liver. We also questioned if certain molecular events were activated at different stages. Therefore, we performed GSEA using differential transcriptomes comparing MASL, MASH or cirrhosis to healthy livers and obtained top shared upregulated functions at a higher resolution. The activation of most immune and cellular anomalies began at MASL. Notably, certain functions, including the innate immune response (*i.e.* neutrophilic response and antigen presentation), cellular damage and rejuvenation (*i.e.* apoptosis and angiogenesis), lipid metabolism and most cytokine and chemokine-mediated signaling were considerably more activated at MASL compared to other activities like adaptive immune response, type II interferon signaling, mechano-signaling and fibrogenesis which were more prominent at MASH (**Figure 2C**). According to the predicted cellular landscape from bulk RNAseq deconvolution, intrahepatic infiltration of innate immune cells preceded that of adaptive immune cells (**Figure 2D-E**). Importantly, the expression patterns of the DEGs and the enriched functions exhibited a gradual up- or downward trend, instead of a stage-specific up- or downregulation, as MASLD advanced (**Figure 2B-C**). This observation highlights a progressive and evolving transcriptomic shift in response to prolonged exposure to MASLD-associated stressors, despite distinct histologic manifestations between different stages.

The onset of MASLD exhibits sexual dimorphism, with females having lower risk for the disease at reproductive age but the protective effect starts to wear off after menopause^40^. Given that gender was a strong covariate in the liver transcriptomes (**Figure 2A**), we investigated the interaction term with gender and MASLD stages to identify hepatic genes that displayed diverse expression pattern between males and females as the disease advanced. There were 83 DEGs showing significant gender-disease stage interaction and a uniform expression trend compared to healthy livers (**Figure S1C, Supplementary Data 5**). Three Y-chromosome genes, namely *EIF1AY*, *KDM5D* and *USP9Y*, which were markedly overexpressed in male livers, could potentially drive the gender-dependent transcriptomic differences during MASLD (**Figure S1C-D**). According to predicted gene-gene interaction, they were responsible for translation initiation, and interacted with proteins bearing ankyrin repeat domain or involved in mRNA processing and cell cycling, highlighting a transcriptional regulatory role which may determine cell fate (**Figure S1E**). Nonetheless, it is noteworthy that the number of genes showing gender-disease stage interaction was relatively few and most DEGs driving the immunopathologies and cellular injury of MASLD were unaffected by gender. Therefore, the intrinsic transcriptomic differences between male and female livers may have limited influence on the sexual dimorphism in MASLD progression. Once MASLD has developed, this gender effect becomes marginal. In short, several Y-chromosome genes may contribute to the sex differences in MASLD, but their actual effect size should be validated.

### Unsupervised clustering predicts risk of progressive MASLD

Given the limited gender influence directly on DEG related to disease progression, there was considerable interest in curating gene signatures for MASLD staging. To shortlist key genes for transcriptomic-guided staging while minimizing computational resources and time demands, we applied the Boruta feature selection algorithm to the 820 DEGs between MASLD stages^31^. The analysis yielded 66 key genes, of which 56 were progressively upregulated as MASLD deteriorated (collectively known as “MASLD gene signature”) (**Figure 3A**). To broaden the utility of the gene signature, we performed an inter-species validation using the liver transcriptomes from our in-house diet-induced MASLD mouse model^33^. We refined the gene signature by selecting genes with a mouse ortholog that showed a consistent expression pattern with the human counterpart. This process resulted in a 39-gene human-mouse harmonized MASLD signature (**Supplementary Data 3**). This curated gene set is useful for quantification and relative comparison of MASLD severity in mice via a ssGSEA-based approach as described in our recent study^33^ (**Figure S2A-B**). An end-to-end analytical application using liver transcriptomes for MASLD staging is also available in our MegaMASLD webtool.

**Figure 3:**
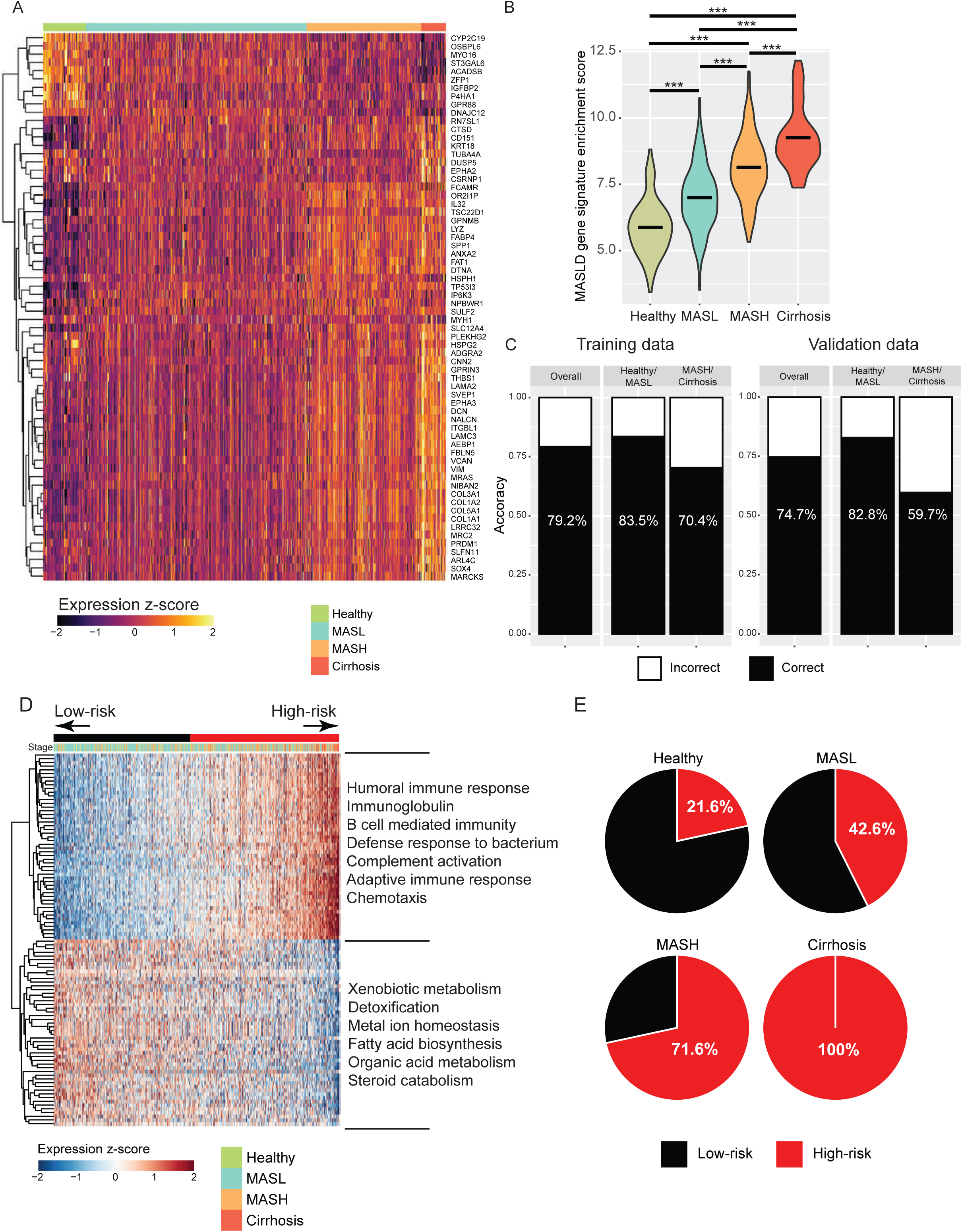
Consensus non-negative matrix factorization (cNMF)-based unsupervised clustering predicts risk of progressive MASLD. **A.** Heatmap of 66 MASLD gene signature based on Boruta feature selection. **B.** Enrichment scores of the human-mouse harmonized MASLD gene signature of liver transcriptomes from varying stages. Crossbars indicate mean scores. *p<0.05, **p<0.01, ***p<0.001 (pairwise t test with Benjamini-Hochberg adjustment). **C.** Accuracy of dichotomous patient classification into early (healthy and MASL) or advanced (MASH and cirrhosis) in training (left) and validation (right) datasets based on a logistic regression that models the relationship between enrichment scores of the human-mouse harmonized MASLD gene signature and the binary stages. **D.** Heatmap of the top 50-gene signatures from cNMF-based unsupervised clustering of the liver transcriptomes. The patients are stratified into two subgroups, defined as low- and high-risk subgroups for progressive MASLD, respectively. **E.** Percentage of patients who have high risk for progressive MASLD separated by their histologic staging. \**P* < 0.05, ** *P* < 0.01, and *** *P* < 0.001 (pairwise t test with Benjamini-Hochberg adjustment).

When applied on human liver transcriptomes, the analysis revealed a significant positive association between the enrichment scores and histologic staging. This finding underscored a potential use of transcriptomic data for patient stratification on a continuous scale, rather than relying on discrete ordinal scoring between groups (**Figure 3B**). To evaluate the diagnostic value of the harmonized gene signature, we trained an ordinal logistic regression model to predict histologic staging based on the enrichment scores. The performance of the model was suboptimal, with overall accuracy of 64.1% and 57.4% using training and validation data, respectively (**Figure S2C**). Intriguingly, when we dichotomized the transcriptomes into early (healthy and MASL) or advanced (MASH and cirrhosis) and modelled the relationship of the enrichment scores and the binary stages using a logistic regression, the predictive power improved to an overall accuracy of 79.2% and 74.7% in training and validation data, respectively (**Figure 3C, S2D**). These observations indicated a correlation between histologic staging and transcriptomic shifts during MASLD progression, although the two did not fully mirror each other. This discrepancy may stem from several factors. Firstly, histological classification is inherently imperfect and prone to inter- and intra-observer variability^41^. Secondly, the heterogenous nature of MASLD may complicate the correlation, as the disease encompasses a wide spectrum exhibiting varying immunopathologies and potentially orchestrated via distinct underlying mechanisms. As shown in our analysis, the gene expression alterations underpinning MASLD are dynamic and cumulative (**Figure 2B-C**). Such quantitative measure of gene expression may even offer advantages over discrete and observer-dependent histologic classifications, providing a more robust risk stratification strategy for MASLD patients.

To leverage the unique strengths of transcriptomic data, we performed an unsupervised clustering using cNMF which identified patient groupings solely on their liver transcriptomic profiles. As the clustering strategy assumed no prior knowledge, it could reveal hidden patterns or subgroups with different risk profiles. The integrated liver dataset was stably clustered into two subgroups, with the highest silhouette value (>0.8) and excellent cophenetic correlation coefficient (>0.95) (**Figure S3A-B**). Using top 50-gene signatures from each subgroup, we showed that one of the subgroups was over-represented by healthy or MASL patients while the other subgroup had more advanced MASLD patients (*i.e.* MASH and cirrhosis). Consequently, these patients were stratified into low- and high-risk, respectively (**Figure 3D, Supplementary Data 6**). Notably, the proportion of high-risk patients almost doubled from 21.6% healthy individuals to 42.6% in MASL patients. The prevalence increased further to 71.6% in MASH and 100% in cirrhotic patients (**Figure 3E**), highlighting the potential usage of this strategy to predict patients at risk of progressive MASLD, especially MASL-to-MASH transition. This analysis demonstrated the capability of transcriptomic analysis to predict risk of disease progression, offering a promising approach for improving patient outcomes through precision medicine.

### Dysregulated humoral immune response underpins MASL-to-MASH transition

Based on our analysis, more than 2 in 5 MASL patients were considered high-risk for progressive MASLD (**Figure 3E**). PCA of their transcriptomic profiles supports the risk stratification despite the shared histologic diagnosis (**Figure 4A**). Demographically, high-risk MASL patients were older in both male (high-risk: 56.2 years; low-risk 49.6 years) and female patients (high-risk: 58.7 years; low-risk: 51.5 years) and marginally more prevalent in male (46.3% in males, 38% in females, p=0.076).

**Figure 4:**
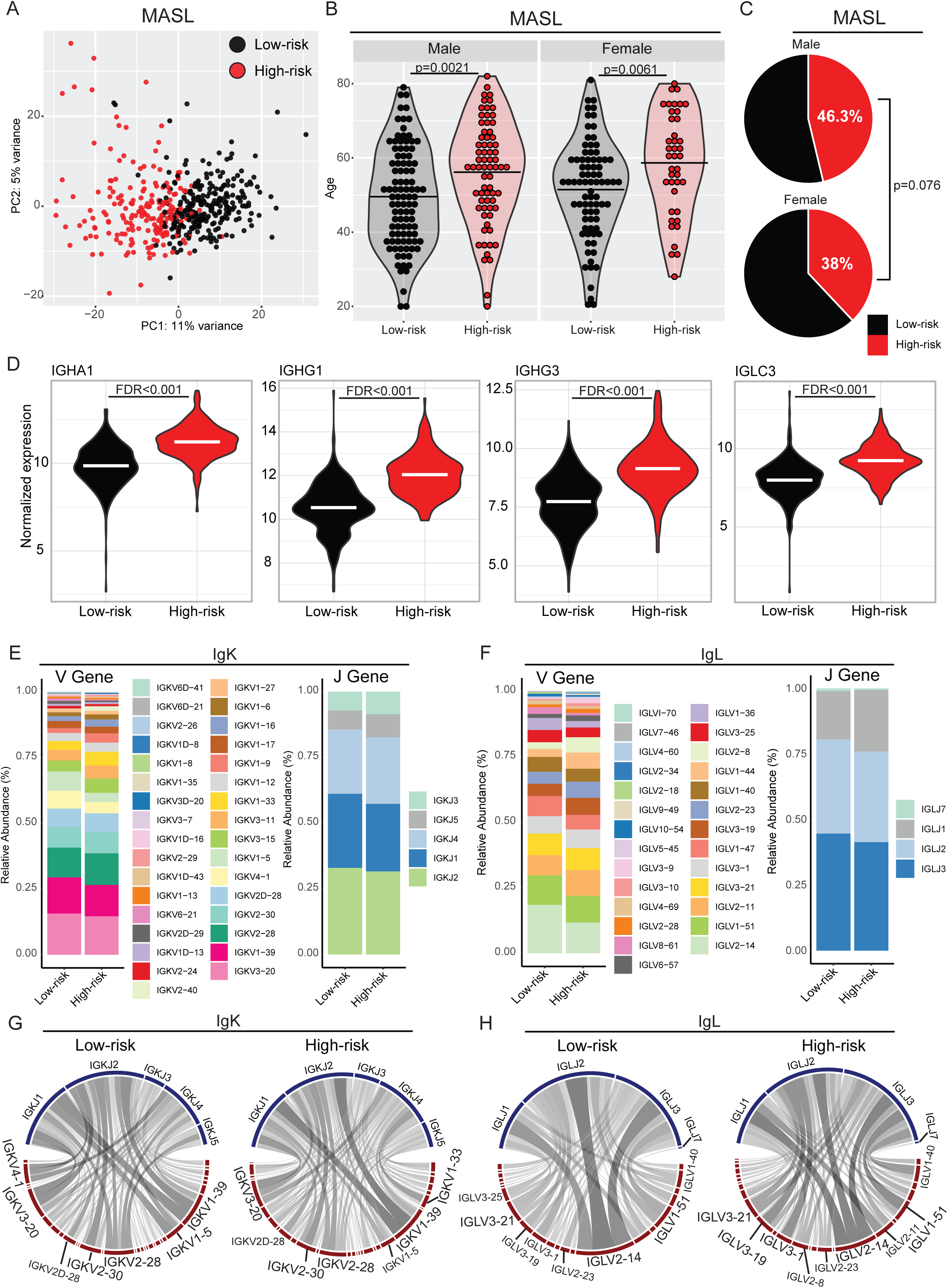
Dysregulated humoral immune response underpins MASL-to-MASH transition. **A.** PCA of liver transcriptomes showing distinct clusters based on the two subgroups of cNMF stratification among the MASL patients. **B.** Age distribution between low- and high-risk subgroups among MASL patients. Bars in the density plots indicate mean age. **C.** Percentage of high-risk patients for progressive MASLD between male and female MASL patients. Crossbars indicate mean expressions. **D.** Expression of immunoglobulin genes between low- and high-risk subgroups among MASL patients. **E-H.** V- and J-gene usage (**E-F**) and VJ recombination of top 3 clonotypes (**G-H**) in κ- (IgK) and λ- (IgL) light chain immunoglobulins between low- and high-risk subgroups among MASL patients.

In clinical settings, close monitoring and timely management are warranted for these at-risk patients who are prone to MASL-to-MASH transition. Hence, there is a critical need to identify reliable markers for patient stratification. Our findings indicate that the gene signature-associated with the high-risk subgroup were closely linked to dysregulation of the humoral immune response. This is evidenced by the overexpression of immunoglobulins, including IgA, heavy and light chains of IgG (**Figure 3D, 4D**). Likewise, the upregulation of these immunoglobulin genes in fibrotic and cirrhotic livers was also found using another liver database – GepLiver^19^ (**Figure S4A**). Assessment of the diversity of the BCR repertoire at the CDR3 regions revealed significantly higher Gini Coefficient of *IGL* (λ-light chain immunoglobulin), but not *IGH* and *IGK* (heavy chain and κ-light chain immunoglobulins), in the high-risk subgroup among MASL patients (**Figure S4B**). The gene usage of VDJ and constant regions of *IGH* were comparable between low- and high-risk subgroups (**Figure S4C-F**). Interestingly, changes in VJ recombination of top three most abundant clonotypes of *IGH* were observed. In low-risk subgroup, IGHV3-23-IGHJ4 was the predominant combination while in high-risk subgroup, combinations such as IGHV3-30-IGHJ4, IGHV3-49-IGHJ4, IGHV3-53-IGHJ4, and IGHV4-31-IGHJ6 became more abundant **(Figure S4G**). Compared to *IGH*, changes in the V gene usage of *IGK and IGL* were more prominent, as evidenced by the increased abundance of *IGKV1-9*, *IGKV1-16*, *IGKV2-40*, *IGLV1-44* and *IGLV2-8* (**Figure 4E-F**). Likewise, VJ recombination of top three most abundant clonotypes of *IGK* and *IGL* also differed between the two risk subgroups. In *IGK* of high-risk MASL patients, the combinations involving *IGKV1-39* were the dominant clonotypes (**Figure 4G**). Conversely, the *VJ* recombination of *IGL* in high-risk subgroup were more diverse compared to low-risk subgroup (**Figure 4H**).

Together, our results underscored the profound implications of B cell activity and humoral immune response in progressive MASLD. Patients at risk for MASL-to-MASH transition had higher expression of immunoglobulin genes, altered gene usage of light-chain immunoglobulins and distinct VJ recombination, collectively pointing towards a perturbed antibody repertoire that is useful for diagnosis and detection of this at-risk population. Further investigation should explore the biological implications of B cell dysregulation in MASLD progression.

### Pseudotemporal expression of hepatic genes confirms dysregulation of humoral immune response and β-chemokine expression preceding MASL-to-MASH transition

Given a sufficiently large sample size and robust data integration, the liver transcriptomes derived from varying disease stages can be ordered in a pseudo-temporal manner to represent the full MASLD spectrum, despite their diverse origins (**Figure 5A**). This analysis provides a valuable avenue to infer crucial transcriptomic events along the course of MASLD deterioration, without the need for repeated liver biopsies. By applying Slingshot^39^ onto the integrated liver transcriptomes, we mapped out a pseudotime projection that started from predominantly healthy livers and progressed through MASL and MASH, culminating in severe cirrhosis (**Figure 5B-C**). Although the discrete histologic stages were not clearly demarcated by pseudotime inference, given the significant overlaps between samples histologically classified into different severity, the pseudotime trajectory was in better agreement with the cNMF risk stratification. Patients at-risk for progressive MASLD had higher pseudotime scores, and vice versa (**Figure 5C**).

**Figure 5:**
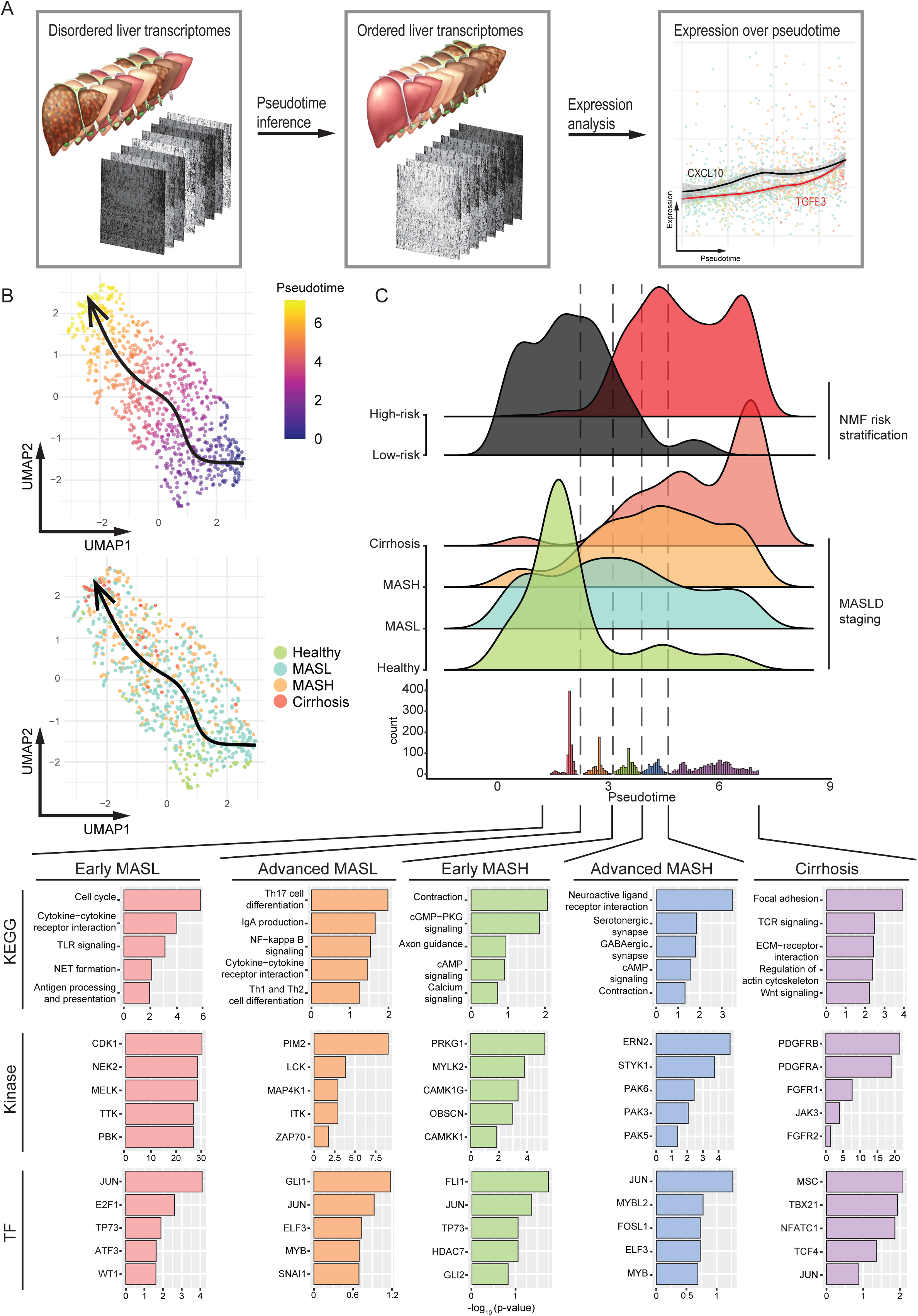
Pseudotemporal analysis of bulk liver transcriptomes reveals new targetable regulomes at various MASLD stages. **A.** Schematic diagram of the pseudotime analysis of bulk liver transcriptomes from MASLD patients with varying disease severity. **B.** Inference of a pseudotime trajectory based on 783 liver transcriptomes. **C.** Distribution of the pseudotime scores of each transcriptome and take-off points of upregulated genes along the pseudotime trajectory. The genes are separated into 5 groups based on the penta-phasic distribution of their take-off points. Top enriched KEGG pathways, protein kinases and transcription factors for each gene group are shown to reveal the key transcriptomic events associated with each transitory stage.

Using the pseudo-temporally ordered data, we computed a take-off point, defined as a (pseudo)time point when the expression of a gene begins to exhibit a significant and continuous upward trend, for every gene. We found 3804 genes with a take-off point (**Supplementary Data 7**). Interestingly, the take-off points of these genes showed a penta-phasic distribution along the pseudotime trajectory, with each peak preceding certain transitory stages, which we broadly categorized into early and advanced MASL, early and advanced MASH, and cirrhosis (**Figure 5C**). Genes with a take-off point at early MASL were associated with cell cycle regulation and innate immune response and were regulated by protein kinases like CDK1, NEK2, MELK, TTK and PBK, suggesting an enhanced cellular proliferation to overcome hepatic injury. As the disease progressed into advanced MASL, there were more severe inflammatory and adaptive immune response as genes related to NF-κB signaling, CD4 T cells and B cells starts to spike. These genes were likely responsible for driving the MASL-to-MASH transition. During early MASH, genes regulated by protein kinase cGMP-dependent 1 (PRKG1) and Ca²⁺/calmodulin-dependent protein kinases were highly expressed (**Figure S5A**). These events coincided with overexpressed of contraction-related genes, pointing to a calcium and cGMP-mediated transformation of quiescent hepatic stellate cells into their activated form to fuel fibrogenesis. The heightened fibrogenic activities of hepatic stellate cells continued in advanced MASH as evidenced by overrepresentation of genes related to serotonergic, GABAergic and P21-activated kinases (PAK) signaling pathways. In cirrhotic stage, the pseudotime analysis predicted increased PDGFR and FGFR signaling which may underpin further fibrogenic activity or malignant transformation into hepatocellular carcinoma (**Figure S5B**). Based other liver single cell atlases, hepatic stellate cells exhibit high expression levels of *PRKG1*, *FGFR1*, *FGFR2*, *PDGFRA* and *PDGFRB* (**Figure S6**). Therefore, inhibitors of these molecular switches could serve as potential anti-fibrotic modalities against liver fibrosis. Additionally, c-JUN, a key component of the activator protein-1 (AP-1) transcription factor complex, was implicated in all phases of MASLD pathogenesis (**Figure 5C, S5C**). It is ubiquitously expressed in most cell types in the liver environment, highlighting a crucial role of c-JUN-mediated regulatory network in immune, stromal and parenchymal alterations during MASLD progression (**Figure S6**).

Furthermore, we examined the pseudotemporal expression of heavy and light chain immunoglobulin genes. The take-off points of most immunoglobulin genes preceded MASL-to-MASH transition based on cNMF risk stratification, in line with altered intrahepatic BCR repertoires in patients at-risk for progressive MASLD (**Figure 6A**). The surge in immunoglobulin gene expression occurred in tandem with the upregulation of C-C chemokines/β-chemokines like *CCL2, CCL3* and *CCL18* but showed less consistency with the expression profiles of α-chemokines and interleukins (**Figure 6B, Supplementary Data 7**). Hence, β-chemokine-mediated infiltration of B-lymphocytes into the liver may bolster a critical event in orchestrating the advancement of MASL to MASH.

**Figure 6:**
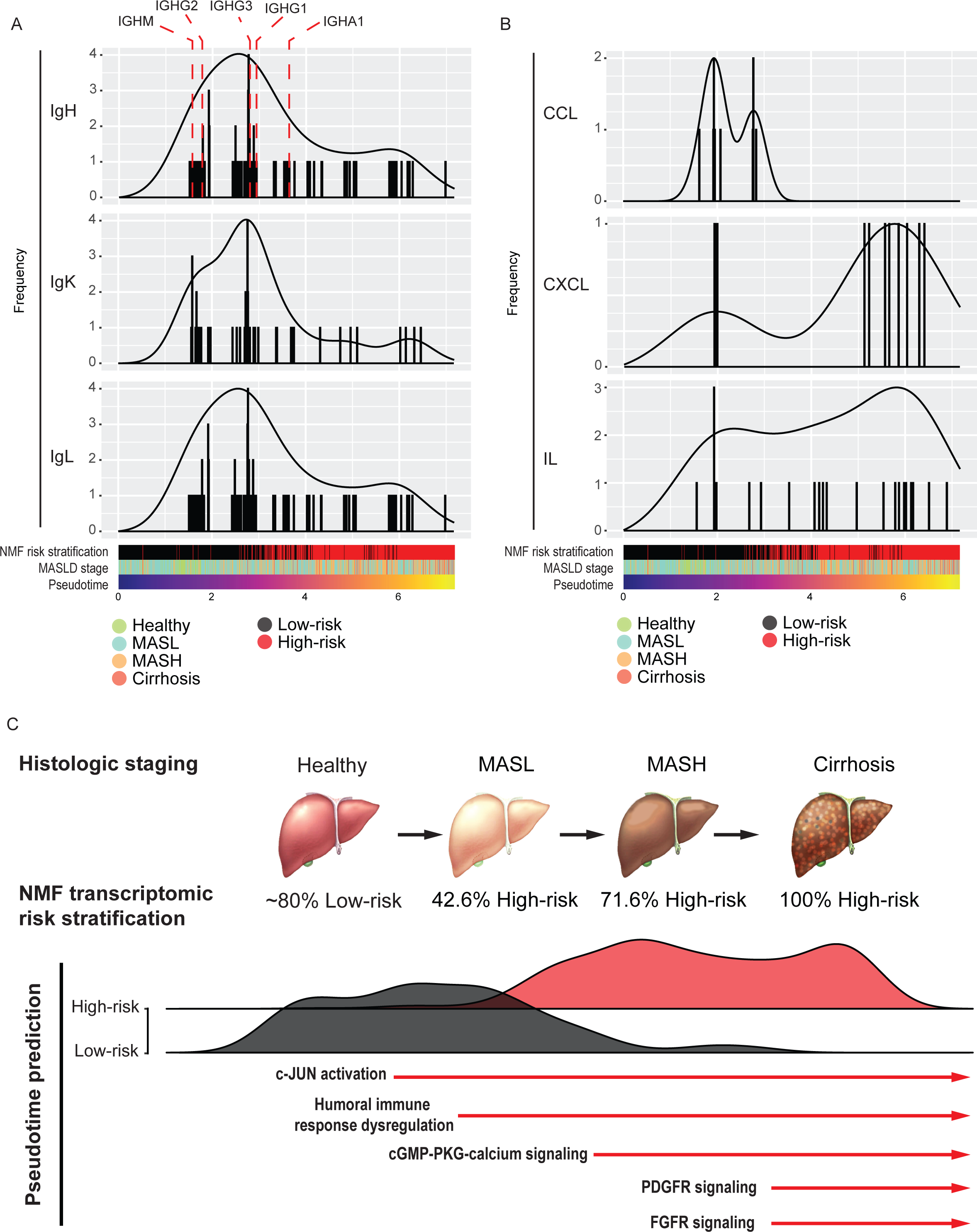
Abnormal surge in immunoglobulin and β-chemokine expression as precedents to the MASL-to-MASH transition. **A-B.** Distribution of the take-off points of heavy and light chain immunoglobulins (IgH, IgK and IgL; **A**) and signaling molecules like β- (CCL) and α-chemokines (CXCL) and interleukins (IL) along the pseudotime trajectory. **C.** Summary of the key findings based on the integrative transcriptomic analysis of patient MASLD livers.

In short, the pseudotime analysis not only reaffirmed the importance of dysregulated humoral immune response in MASLD pathogenesis, but also pinpointed a several signaling regulomes, namely c-Jun, cGMP-PKG-calcium signaling, PDGFR and FGFR signaling which can serve as exploitable targets to curtail key molecular events and slow down the progression of the liver disease (**Figure 6C**). This work highlights the potential of pseudotemporal transcriptomic analysis in uncovering the molecular mechanisms driving MASLD progression and provides valuables insights for developing targeting therapeutic strategies, ultimately advancing precision medicine for MASLD patients.

## Discussion

Here, we present a robust integrative analysis of a large MASLD liver transcriptome dataset comprising 830 patients from 16 independent cohorts. The analysis reveals a progressive and evolving transcriptomic shift as MASLD advances. Unsupervised clustering of the continuous gene expression data categorizes individuals into low- and high-risk subgroups for progressive MASLD. The high-risk subgroup, predominantly older males, is more prone to MASL-to-MASH transition and exhibits a perturbed humoral immune response, characterized by higher immunoglobulin gene expression and altered antibody repertoires compared to low-risk subgroup. Pseudotime analysis of the integrated hepatic transcriptomes supports the disruption of B-cell activity during MASL-to-MASH transition, and predicts druggable targets, including c-Jun, PRKG1, Ca²⁺/calmodulin-dependent protein kinases, and PDGFR and FGFR signaling. The integrated dataset and analytical utilities are available at MegaMASLD (https://bioanalytics-hs.shinyapps.io/MegaMASLD/), an interactive R Shiny app encompassing over 800 human MASLD liver data encompassing the full MASLD disease spectrum. By focusing specifically on the disease context of MASLD, MegaMASLD provides unique insights on molecular mechanisms, disease progression, and potential therapeutic targets, which are under-represented by other liver single-cell atlases, such as GepLiver^19^, Liver Cell Atlas^42^, liver cirrhosis^43^ and liver plasticity^20^ landscapes. Hence, MegaMASLD is a great complement to existing liver databases, allowing researchers and clinicians to delve further in the disease context for advancing MASLD research and clinical applications.

MASLD is associated with significant alterations in hepatic transcriptomes linked to inflammation, immune response, hepatic injury, and fibrogenesis^11,15,44^. While various studies have curated gene signatures based on the histological staging to predict disease severity, our analysis underscores pronounced inter-study inconsistencies. For instance, Ryaboshapkina & Hammer (2017) identified a 218-gene MASLD progression signature, while Hasin-Brumshtein et al. (2022) reported a 130-gene signature comparing MASH versus healthy/MASL livers^11,44^. Notably, only 25 genes are shared between the two major gene sets, and merely four overlapping with our human-mouse harmonized MASLD gene signature (*FABP4*, *FAT1*, *SPP1*, and *SULF2*). Furthermore, while these gene signatures are useful for determining the relative disease severity between samples, their accuracy to predict the histologic stages is considerably low. This highlights the variability in transcriptomic-guided staging versus histological assessment, which is likely due to heterogeneity of the nature of MASLD and inter- and intra-observer variability in histopathological assessment^41^. Our study clearly showed that the transcriptomic changes in MASLD is cumulative and dynamic. The quantitative gene expression changes may offer more granularity and finer resolution than discrete histologic classifications in risk stratification of MASLD patients.

Our study offers two histologic-independent strategies, cNMF unsupervised clustering and pseudotime analysis, revealing novel insights into the pathogenesis of MASLD. Unsupervised clustering predicts two subgroups based on the risk for advanced MASLD, identifying over 40% of MASL individuals as high-risk for MASL-to-MASH transition or progressive MASLD. Pseudotime analysis, another histologic-independent analysis, elucidates dynamic transcriptomic changes during MASLD progression, supporting alterations in BCR repertoire and the role of humoral immune response in MASLD pathogenesis. These at-risk patients exhibit higher expression of heavy-chain immunoglobulins and varying gene usage of light-chain immunoglobulins κ and λ. Altered antibody repertoires may arise from aberrant deposition and maturation of intrahepatic B cells into plasma cells during the onset of steatohepatitis^45,46^. Additionally, a subset of MASLD patients have elevated levels of auto-reactive antibodies, which may increase the risk of hepatic damage and fibrosis^47,48^. Given the lack of reliable screening methods to identify MASL patients at risk for developing MASH in clinical settings, dysregulated antibody repertoires can serve as potential markers to stratify these high-risk patients who may require more proactive management. Further investigation is needed to dissect the exact role of humoral immune response in MASLD pathogenesis.

Pseudotime analysis also identifies overexpressed β-chemokines coinciding with the onset of dysregulated humoral immune response, with multiple C-C motif chemokines (CCLs) being upregulated. Certain CCLs, such as CCL3 and CCL19, play vital role in the positioning and homing of B cells to form germinal centers^49,50^. Given the complex signaling cascades mediated by different CCLs, targeting individual CCLs or their receptors (CCRs) may not effectively attenuate MASH deterioration. For instance, Cenicriviroc, a dual CCR2/CCR5 antagonist, has shown limited success in resolving liver fibrosis in a phase III clinical trial, highlighting the need for therapeutic strategies that block a wide range of CCL-CCR interaction^51^.

Additionally, pseudotime analysis identifies other targetable kinase signaling cascades and transcription factors governing the key molecular events at different phases of MASLD. Some of these targets are well-established, while others are potentially novel, presenting new avenues for therapeutic intervention. For example, we found that the regulome of c-Jun are involved in all phases of MASLD pathogenesis. This aligns with previous studies showing that the ablation of hepatocyte-specific c-Jun expression effectively prevented diet-induced liver fibrosis and hepatocellular carcinoma in mice^52,53^. Conversely, the analysis predicts several transcriptomic reprogramming networks linked to the activation of hepatic stellate cells, including the signaling pathways mediated by FGFR, PDGFR, PAK, PRKG1 and CAMK kinase families. While the therapeutic potential of these novel target remains largely unclear, apart from selected preclinical studies^54,55^, many pre-existing kinase inhibitors are available for repurposing to validate their anti-fibrotic effect in MASLD.

While our study has brought forth key molecular events in hepatic transcriptomes that underpin MASLD pathogenesis, the lack of comprehensive patient metadata, such as ethnicity, lifestyle factors and menopausal status, limit the analysis. Other health related measurements like body-mass index and lipid profiles can facilitate the identification of minor disease subtypes, *e.g.* lean MASLD, to reveal potentially unique markers and pathogenesis. Our findings are derived from human liver RNAseq data, experimental validation using preclinical models are highly recommended to confirm the therapeutic effect of the potential drug targets.

## Conclusion

In summary, we have integrated diverse transcriptomes from healthy, MASL, MASH and cirrhotic livers, revealing progressive transcriptomic shifts not perfectly aligned with histologic staging where distinct microscopic features are observed. Our analysis highlights significant dysregulation of the humoral immune response during MASL-to-MASH transition and offer tools for risk stratification and drug target prediction. These applications, available on a user-friendly web platform, MegaMASLD, bridges large omics datasets and clinical utility, aiming to improve the precision medicine of MASLD. It enhances the accessibility of complex transcriptomic data and empowers researchers and clinicians to translate molecular insights into tangible health benefits.

## Acknowledgements

H.S. Cheng is a recipient of LKCMedicine Dean’s Postdoctoral Fellowship from Nanyang Technological University Singapore. D. Chua is a recipient of the Nanyang Presidential Graduate scholarship. Part of the computational work was performed using resources from the National Supercomputing Centre (NSCC), Singapore.

