## Supplementary figures and images for "MegaMASLD: An interactive platform for exploring stratified transcriptomic signatures in MASLD progression"

### Figure S1

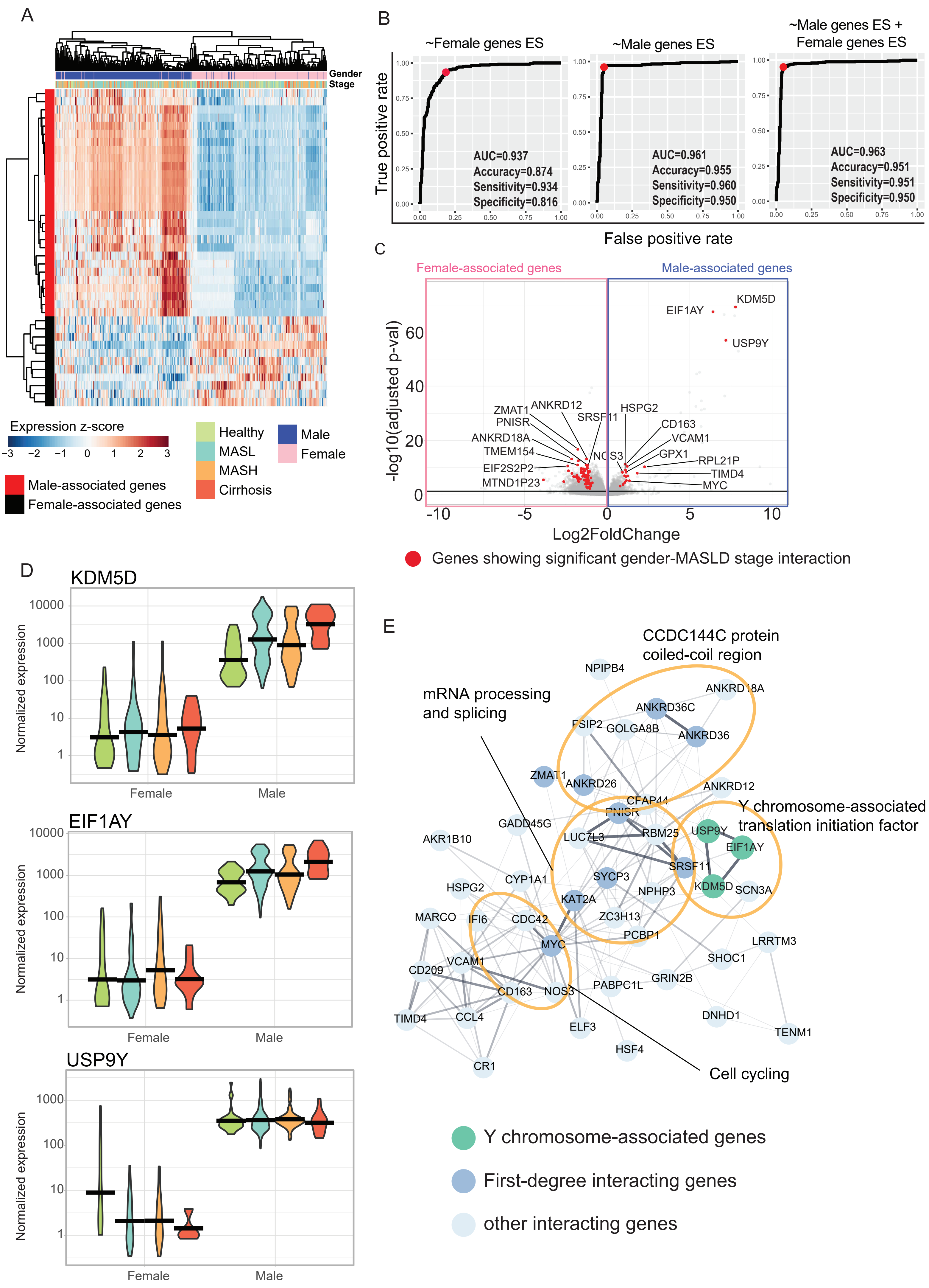

### Figure S2

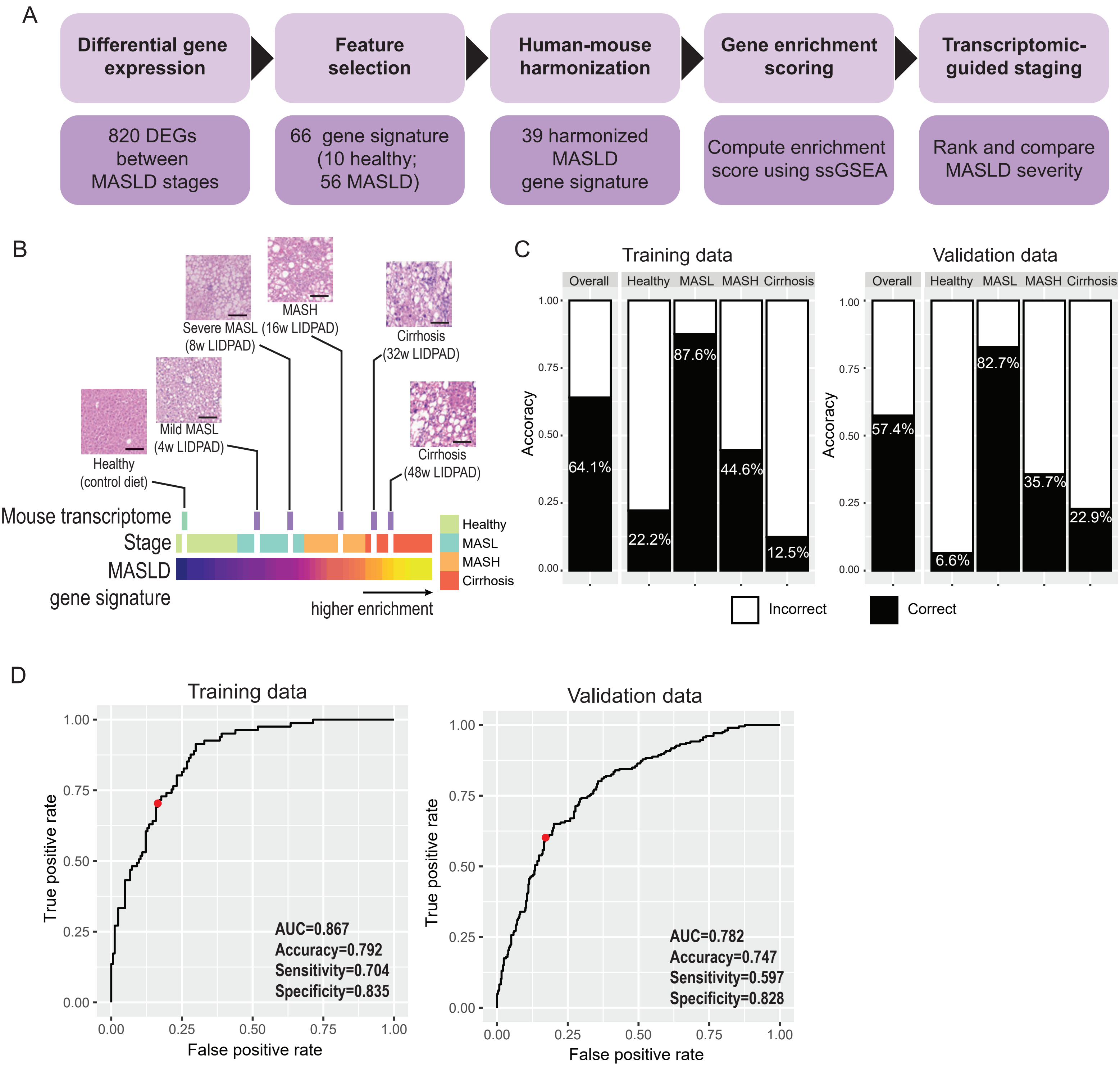

### Figure S3

Cheng et al. Figure S3

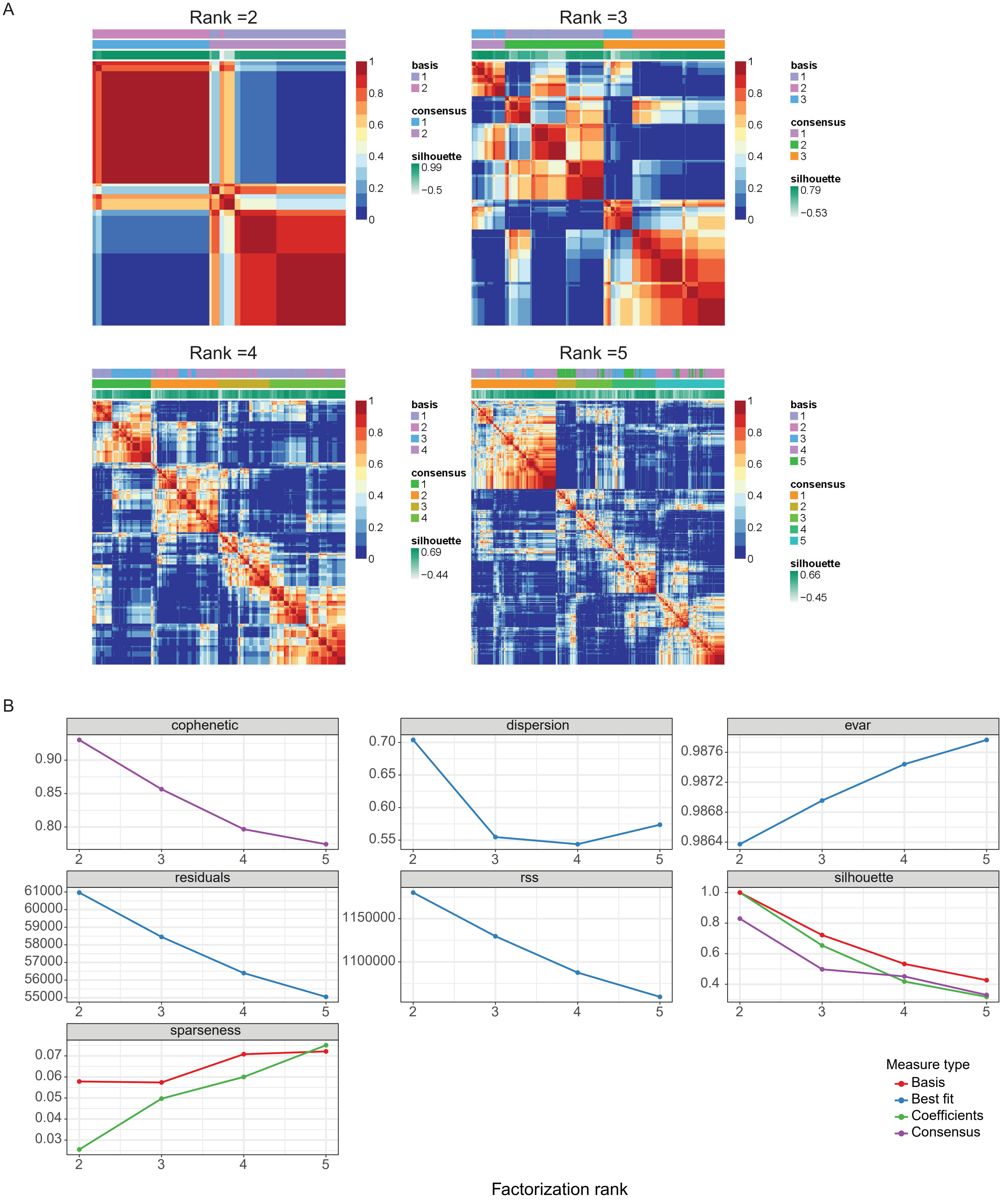

### Figure S4

Cheng et al. Figure S4

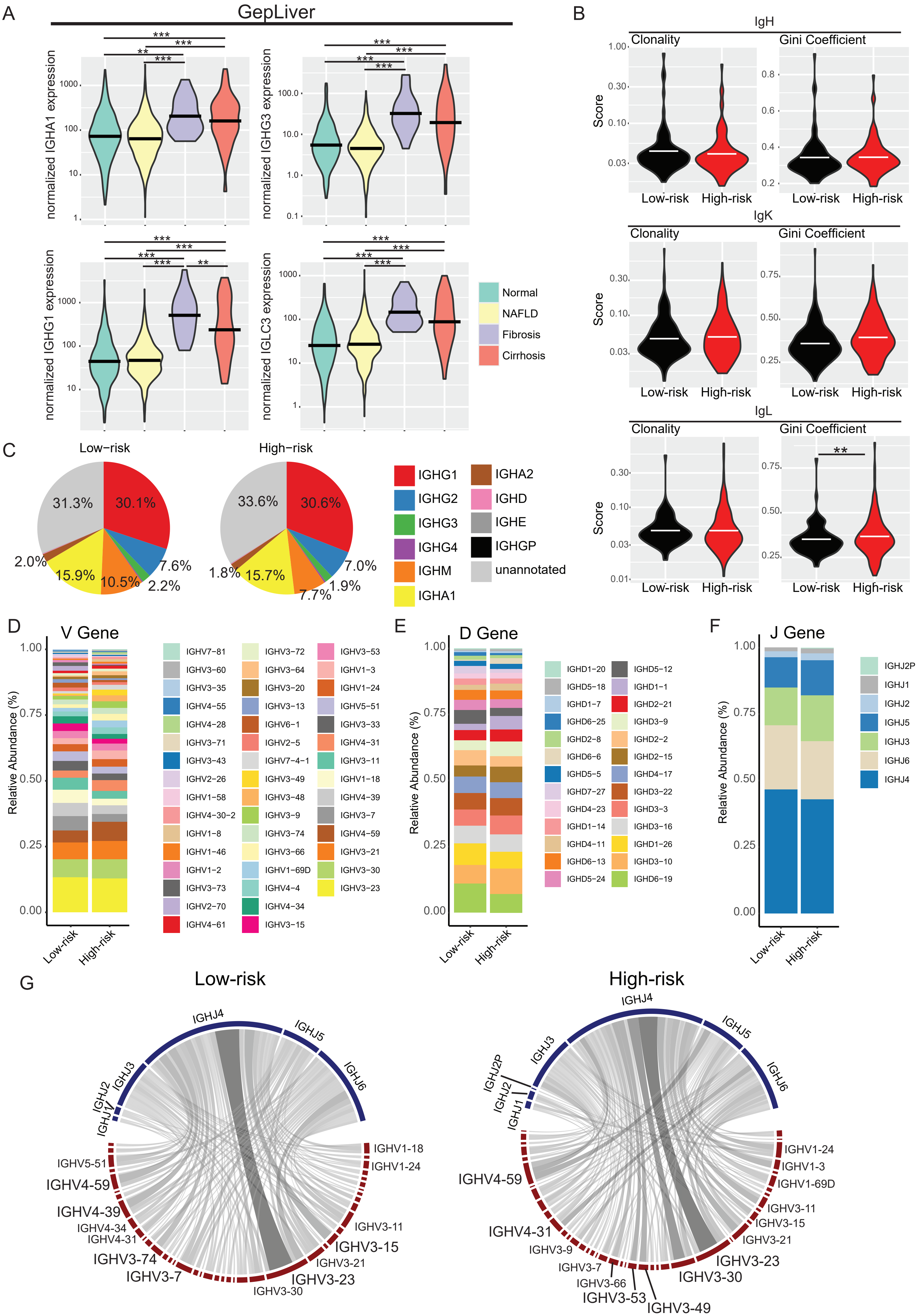

### Figure S5

Cheng et al. Figure S5

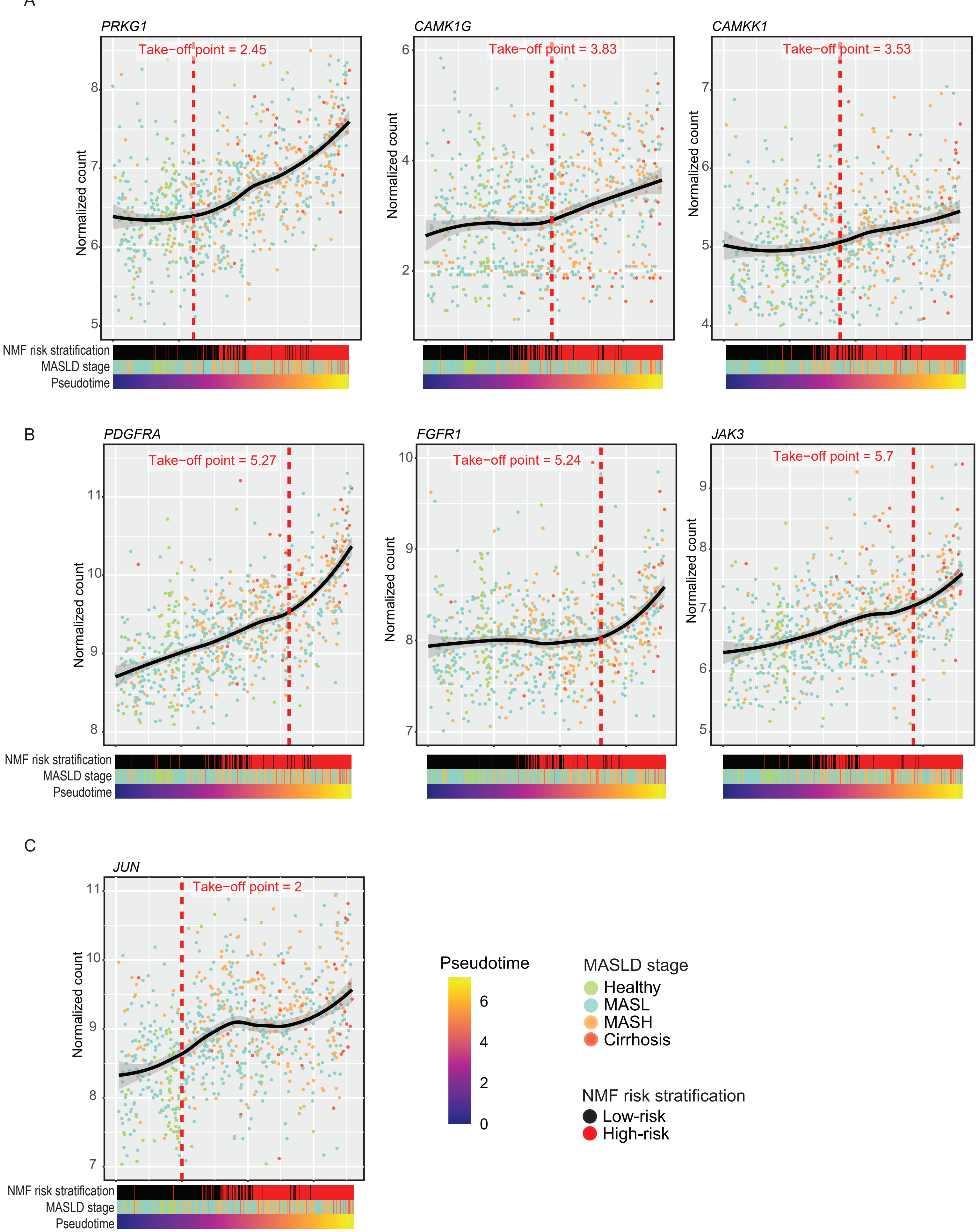

### Figure S6

Cheng et al. Figure S6

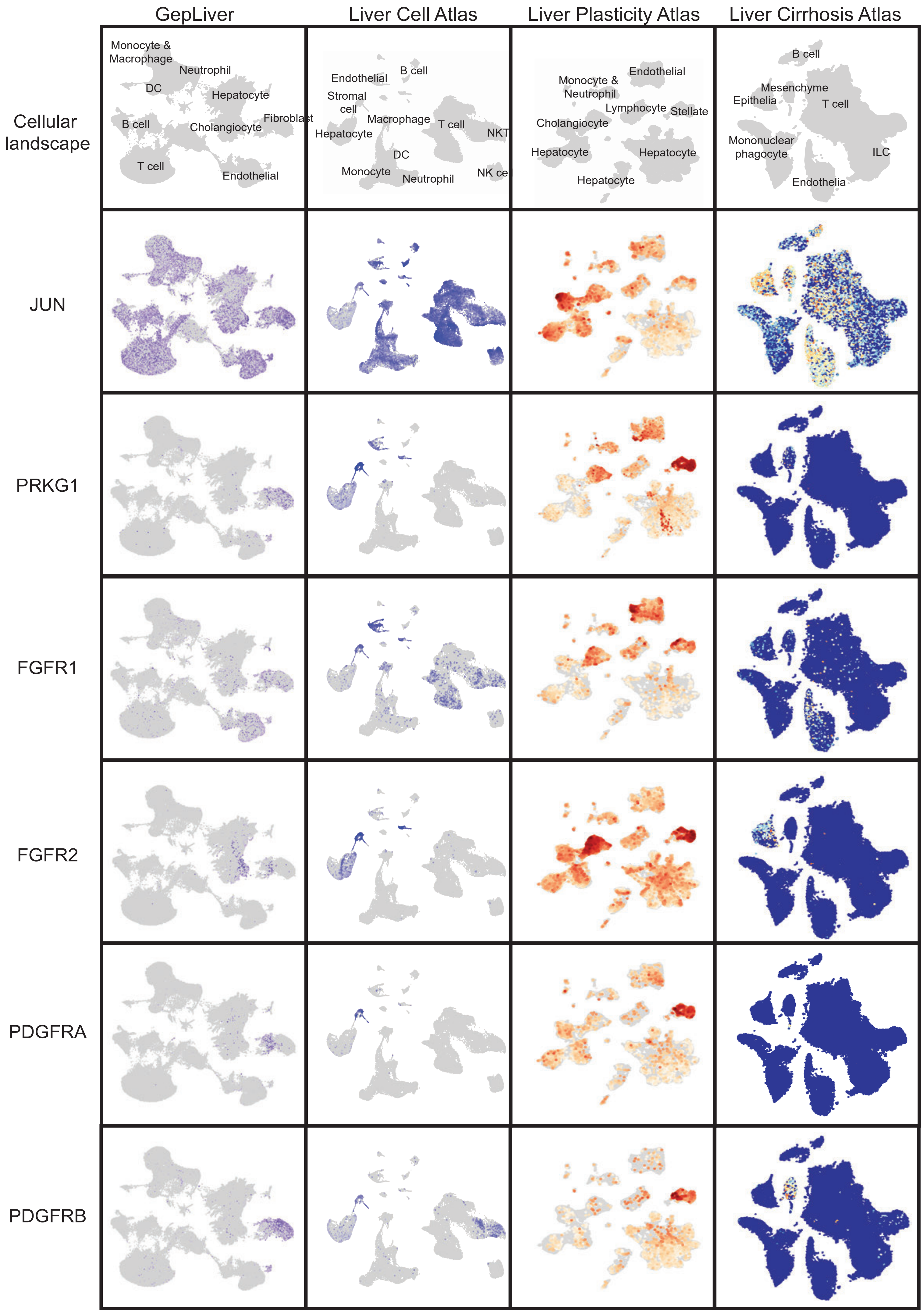
