## Supplementary information for "MegaMASLD: An interactive platform for exploring stratified transcriptomic signatures in MASLD progression"

### MegaMASLD: An interactive webtool revealing transcriptomic shifts and key molecular events in human MASLD progression

Running title: Molecular insights into MASLD pathogenesis

*Correspondence:

**H.S.C.**, Lee Kong Chian School of Medicine, Nanyang Technological University Singapore, 11 Mandalay Road, Singapore 308232.

#### Supplementary Table

**Table S1: Thresholds of liver histology scores for MASLD staging.**

| MASLD stage | Criteria |
|---|---|
| Healthy | NAS=0 & Fibrosis=0 |
| MASL | NAS ≤ 3 & Fibrosis ≤ 2 OR NAS ≤ 4 |
| MASH | NAS > 3 & Fibrosis > 2 OR NAS > 4 |
| Cirrhosis | Fibrosis=4 |

#### Supplementary Files

**Supplementary Data 1**: Metadata of liver transcriptomes

**Supplementary Data 2**: Gender-associated genes

**Supplementary Data 3**: Human-mouse harmonized MASLD gene signature

**Supplementary Data 4**: Differentially expression genes between MASLD stages

**Supplementary Data 5**: DEGs with gender-MASLD stage interaction

**Supplementary Data 6**: Signatures of cNMF clusters

**Supplementary Data 7**: Take-off points from pseudotime analysis

#### Supplementary Figure Legends

**Figure S1. Transcriptomic-guided gender prediction tool and interaction analysis between gender and MASLD stages.**

**A.** Heatmap of 118 gender-specific DEGs between male and female livers.

**B.** Receiver operating characteristic (ROC) analysis of three logistic regressions that model the relationship between enrichment scores of female-specific genes, male-specific genes or both and patient genders.

**C.** Volcano plot of 83 DEGs showing significant interaction effect between gender and MASLD stage.

**D.** Expression of three Y-chromosome genes (*KDM5D*, *EIF1AY* and *USP9Y*) between male and female livers at different MASLD stages. Crossbars indicate mean expressions.

**E.** Gene-gene interactome of 83 DEGs exhibiting significant interaction effect between gender and MASLD stage.

**Figure S2. Establishment of human-mouse harmonized MASLD gene signature for transcriptomic guided staging.**

**A.** Workflow to establish a human-mouse harmonized MASLD gene signature for transcriptomic guided staging.

**B.** Application of the transcriptomic guided staging on liver transcriptomes of LIDPAD mouse model treated for 4-, 8-,16-, 32- and 48 weeks (w). Healthy control mice were treated with a refined matched control diet.

**C.** Accuracy of patient classification into the four histologic stages (healthy, MASL, MASH and cirrhosis) in training (left) and validation (right) datasets based on an ordinal logistic regression model that models the relationship between enrichment scores of the human-mouse harmonized MASLD gene signature and MASLD stages.

**D.** ROC analysis of the ordinal logistic regression model for MASLD staging based on the enrichment scores of the human-mouse harmonized MASLD gene signature.

**Figure S3. Quality assessment of the cNMF-based unsupervised clustering method for risk stratification.**

**A.** Consensus maps from 10 iterations of NMF at rank=2 to 5.

**B.** Rank survey analysis to determine optimal rank. Rank=2 was selected due to the highest silhouette value (>0.8) and excellent cophenetic correlation coefficient (>0.95), suggesting the liver transcriptomes are stably clustered into two subgroups.

**Figure S4. B-cell receptor (BCR) repertoire analysis of low- and high-risk MASL patients for progressive disease.**

**A.** Normalized expression of immunoglobulin genes (*IGHA1, IGHG1, IGHG3* and *IGLC3*) between normal, NAFLD, Fibrosis and Cirrhosis livers obtained from GepLiver database. Crossbars indicate mean scores. *p<0.05, **p<0.01, ***p<0.001 (pairwise t test with Benjamini-Hochberg adjustment)

**B.** Diversity analysis of heavy and light chain immunoglobulins (IgH, IgK and IgL) based on clonality and Gini coefficients. Higher scores of both metrics indicate domination by selected/a single BCR sequence. **$p<0.01$ (student's T test).

**C.** Composition of different heavy-chain immunoglobulin subtypes in low- and high-risk MASL patients for progressive MASLD.

**D-G.** V-, D- and J-gene usage (**C-E**) and VJ recombination of top 3 clonotypes (**F**) in heavy chain immunoglobulin between low- and high-risk subgroups among MASL patients.

**Figure S5. Take-off points of key regulatory genes driving transitory stage based on pseudotemporal analysis.**

**A-C.** Take-off points of *PRKG1*, *CAMK1G* and *CAMKK1* which are linked to MASL-to-MASH transition (**A**), *PDGFRA*, *FGFR1* and *JAK3* which are linked to cirrhotic livers (**B**) and *JUN* which remains overexpressed from early to advanced MASLD.

**Figure S6. Expression of key regulatory genes in other liver single-cell atlases.**

UMAP or tSNE plots derived from four hepatic single-cell atlases, GepLiver, Liver Cell Atlas, Liver Plasticity Atlas and Liver Cirrhosis Atlas, showing the expression of *JUN*, *PRKG1*, *FGFR1*, *FGFR2*, *PDGFRA* and *PRGFRB* in the cellular landscape of a liver environment. DC: dendritic cell, NK: natural killer cell ,NKT: natural killer T cell, ILC: innate lymphoid cells.
